# Long-Term Effects of Adolescent 5F-MDMB-PICA Intravenous self-administration: Neurobehavioral Consequences and medial Prefrontal Cortex Dysfunction in Adult Mice

**DOI:** 10.1101/2025.06.26.661592

**Authors:** Francesca Caria, Avraham Libster, Shane Desfor, Francesca Telese, Gaetano Di Chiara, Maria Antonietta De Luca

## Abstract

**Background:** Synthetic Cannabinoids Receptor Agonists (SCRAs) are the largest group of new psychoactive substances monitored worldwide. 5F-MDMB-PICA is a recent SCRA classified as a potent full agonist at CB1/CB2 receptors able to activate the mesolimbic dopamine (DA) transmission in adolescent but not in adult mice. Here, we have studied its reinforcing effects in adolescent mice and characterized the neurochemical and behavioral effects induced in the same animals in adulthood.

**Methods:** We utilized an intravenous self-administration (IVSA) protocol in adolescent (PND 40-56) CD-1 male mice. In adulthood (PND 66-78), we conducted several behavioral and neurobiological assessments including: Sucrose Preference Test (SPT); Resident Intruder Test (RIT); Olfactory Reactivity Test (ORT); brain microdialysis to quantify DA levels in the medial Prefrontal Cortex (mPFC); and fiber photometry analysis using the GCaMP calcium sensor to monitor excitatory neural dynamics in the mPFC after exposure to an aversive odorant.

**Results:** We found that 5F-MDMB-PICA, administered through IVSA in adolescent mice, produced an inverted U-shaped dose-response curve. The dose of 2.5 μg/kg/25ul elicited behavior consistent with drug seeking. Adult mice exposed to 5F-MDMB-PICA during adolescence exhibited significant behavioral and neurochemical changes in adulthood compared to control mice. These behaviors included increased aggression, reduced social interaction, an anhedonic state, and an abolishment of mPFC DA response to an aversive odorant, as measured by in vivo brain microdialysis. Moreover, fiber photometry analysis of excitatory neuronal activity in the mPFC showed diminished calcium activity in response to the same aversive odorant in 5F-MDMB-PICA-exposed mice compared to controls.

**Conclusions:** Notably, this study is the first to demonstrate that adolescent mice can acquire and sustain IVSA of 5F-MDMB-PICA. Furthermore, it highlights the long-term behavioral and neurochemical changes associated with adolescent exposure to 5F-MDMB-PICA, underscoring the potential detrimental effects of its use during this critical developmental period.

## Introduction

The widespread use of synthetic cannabinoid receptor agonists (SCRAs) has become a major public health concern, driven by their harmful effects and increasing consumption, particularly among adolescents and young adults (1,2). Their psychoactive properties, combined with the mistaken belief that they are safer than natural cannabis, have contributed to their growing appeal. However, the long-term effects of SCRA consumption, especially when used during critical developmental periods such as adolescence, are increasingly raising concerns within the scientific community, and few data are available in this regard. Unlike natural cannabinoids, such as THC, SCRAs exhibit a faster onset of effects and a shorter duration, with varying clinical presentations influenced by factors such as compound specificity, individual susceptibility, and dosage (3–5). Importantly, repeated SCRA exposure has been associated with more severe and enduring adverse effects, including long-term cognitive deficits and mental health disorders (5,6). The emergence of newer generations of SCRAs, pos additional concerns, as these compounds are associated with increased potency and abuse potential compared to earlier variants (7,8).

One of the most alarming aspects of SCRA use is its prevalence among adolescents (9–11) a particularly vulnerable population. Adolescence represents a critical period of brain development, during which the brain, especially the prefrontal cortex (PFC), is highly susceptible to external influences, including psychoactive substances (12,13). Numerous human studies have shown that cannabis use during adolescence is linked to long-term cognitive and behavioral impairments (14), but there is limited understanding of the specific effects of SCRAs on the developing brain. While preclinical studies have demonstrated that SCRAs like JWH-018 can induce enduring behavioral changes in animal models, such as compulsive behaviors and emotional dysregulation (13), little is known about the effects of more potent, newer-generation compounds like 5F-MDMB-PICA. Given the pharmacological profile of these SCRAs, they may pose even more severe and prolonged neurotoxic risks (8,15).

5F-MDMB-PICA has garnered attention due to its widespread use and toxic effects, including cases of acute intoxication and fatalities (7). In 2019, it was the most commonly detected SCRA in the United States, accounting for 31% of the top 10 SCRA reports to NFLIS-Drug 2019 Annual Report (16). Despite its growing popularity among young adults, the specific effects of this compound on brain development during adolescence remain poorly explored. 5F-MDMB-PICA acts as a full agonist at both CB1 (Ki = 1.25 nM; EC50, 5.07 nM) and CB2 (Ki = 0.43 nM; EC50, 1.65 nM) receptors (17), demonstrating hundreds of times greater activity on CB receptors than THC (8,15,18). Its potency surpasses that of JWH-018, with a 129% increase in CB1 receptor activation (19). The potential risks associated with its use, combined with the lack of comprehensive data in the literature, underscore the urgent need for a deeper understanding of its long-term impacts in order to guide public health policies and develop effective prevention strategies.

This study aims to fill this gap by investigating the long-term effects of adolescent voluntary consumption of 5F-MDMB-PICA, focusing on three main areas: (I) establish the reinforcing properties of the substance in animal models, (II) its long-term behavioral consequences, and (III) the neurochemical and neuronal excitability changes in the medial prefrontal cortex (mPFC) in adulthood. To achieve these goals, we employed a well-established intravenous self-administration (IVSA) mouse model (13), allowing us to assess both the acquisition of- and motivation for drug IVSA during adolescence, as well as the long-term behavioral and neurochemical consequences correlated to voluntary consumption. This approach represents a significant advancement by highlighting the grave consequences of consuming such a potent SCRA, offering a deeper understanding of its long-term impact on brain function.

Moreover, in this study, we explored a range of behavioral outcomes, including social and non-social interactions, aggressive and defensive behaviors, and anhedonic emotional states. Special attention was given to behavioral responses to socially challenged stimuli, an area that has been largely neglected in previous SCRA research. Our neurobiological examination focused on the mPFC, a key brain region involved in emotional regulation and cognitive function (20), verifying the impact of adolescent 5F-MDMB-PICA exposure on subsequent activity. In particular, advanced techniques, such as fiber photometry and neurochemical analysis, were used to characterize the mPFC’s response to olfactory aversive stimuli, contributing to a deeper understanding of the molecular mechanisms underlying the observed behavioral changes.

## Materials and methods

*For additional details, please refer to the Supplementary Methods*.

### Animals and housing

Male CD-1 mice, weighing 22-27 g, were purchased from Envigo, Netherlands and employed from adolescence (PND 35) to adulthood (PND 80). At arrival (PND 21), animals were group-housed (3-6 per cage) under standard conditions. Post-surgery for catheter implantation (PND 35-37), individual housing was implemented in specially designed cages to prevent catheter damage. A controlled feeding regimen was established a day before commencing IVSA procedures (PND 39), providing 3-4 g of food pellets daily to maintain stable body weights throughout the experiments. All animal procedures were compliant with the Guidelines for Care and Use of Mammals in Neuroscience and Behavioral Research, adhering to Italian (D. Lgs 26/2014) and European Council Directive (2010/63/UE) standards, and were approved by the Committee for Animal Wellbeing (OPBA) at the University of Cagliari.

For the fiber photometry experiment, male CD-1 mice were purchased from Charles River Laboratory (CD-1® IGS Mouse (Strain 022). All experimental procedures were approved by the institutional animal care and use committee at the University of California, San Diego.

### Drugs and chemicals

5F-MDMB-PICA was purchased from Cayman Chemical Company (Ann Arbor, MI, USA) and dissolved in 0.5% EtOH, 0.5% Tween 80 and 99% saline.

The 2-methyl-2-propanethiol was obtained from Sigma–Aldrich (CAS Number: 75-66-1), ensuring a nominal purity of at least 99% and was diluted 1:10000 in distilled H_2_O.

### Experimental Design

The study was designed to assess the reinforcing properties and abuse potential of 5F-MDMB-PICA during adolescence and its behavioral and neurochemical effects in adulthood. The experimental protocols included:

i. **IVSA studies during adolescence (Experiments I and II)** All the IVSA procedures were carried out as previously described by Margiani et al. (13) and were conducted over the course of two experiments: In Experiment I a dose-response curve for 5F-MDMB-PICA was established. Mice (n=11) were trained to IVSA 5F-MDMB-PICA employing a progressive dosing protocol, where doses were incrementally increased across sessions from 1 μg/kg to 2.5 μg/kg, and finally to 5 μg/kg per 25 μl infusion. The optimal drug intake was observed at a dose of 2.5 μg/kg per infusion. In Experiment II, another cohort of mice (n=38) were trained to self administer 5F-MDMB-PICA at the dose of 2.5 μg/kg/inf across various reinforcement schedules, including Fixed Ratio (FR1, FR3) and a Progressive Ratio (PR) schedule. Additionally, a control group of 25 mice received a Vehicle solution, following the same experimental protocol.
ii. **Behavioral and neurochemical studies at adulthood (Experiment III)** In Experiment III, mice from Experiment II were divided into three groups and randomly assigned to behavioral tests and a neurochemical study conducted after a 10-day drug-free washout period following the last IVSA session. The groups were divided as follows: Group 1 (5F-MDMB-PICA, n=14; Vehicle, n=6) was subjected to the RIT; Group 2 (5F-MDMB-PICA, n=8; Vehicle, n=8) was subjected to the SPT, followed by the ORT after 7 days (5F-MDMB-PICA, n=7; Vehicle, n=5); Group 3 (5F-MDMB-PICA, n=9; Vehicle, n=9) underwent the neurochemical study (in vivo brain microdialysis) to evaluate the mPFC DA fluctuations after exposure to an aversive odorant (see Supplementary Informations for tests details).
iii. **Fiber Photometry Analysis at adulthood (Experiment IV)** In Experiment IV, a fiber photometry experiment was conducted to investigate the impact of adolescent 5F-MDMB-PICA IVSA on mPFC neural activity in response to the aforementioned olfactory aversive stimulus used in Experiment III.

### Data Analysis and Statistics

All data are presented as mean ± SEM. Normal distribution was assessed using the Shapiro-Wilk and Kolmogorov-Smirnov tests. Non-parametric data were analyzed using the Mann-Whitney U-test. For IVSA studies, two-way repeated measures (RM) ANOVA with Sidak’s multiple comparisons was applied, and Geisser-Greenhouse correction was used when sphericity was violated. Unpaired t-tests were used for PR schedule sessions, and one-way ANOVA followed by Tukey’s post hoc test was applied for 5F-MDMB-PICA intake. Behavioral experiments were analyzed using Student’s t-test for normally distributed data, and the Mann-Whitney U-test for non-normal data. Post hoc tests were performed when significant effects or interactions were found, and significance was set at p < 0.05. For neurochemical studies, two-way RM ANOVA followed by Tukey’s multiple comparisons post hoc test was used. Fiber photometry data were analyzed using MATLAB and Python scripts, while DeepLabCut (https://www.nature.com/articles/s41593-018-0209-y) and SimBA (https://www.nature.com/articles/s41593-024-01649-9) were employed for video analysis. Effect sizes (e.g., Cohen’s d, Hedges’ g, Eta squared, Partial Eta squared, R², and correlation coefficient r) were calculated to complement statistical significance and provide practical relevance of the results. Analyses were conducted using GraphPad Prism 8 (Boston, Massachusetts, USA).

## RESULTS

### Reinforcing properties of 5F-MDMB-PICA during adolescence

In an effort to delineate the response pattern of male adolescent CD1 mice, our initial investigation focused on establishing the dose-response curve for 5F-MDMB-PICA within the IVSA experimental framework (PND 40-66). This decision was based on prior research conducted on adult C57 mice (21) and CD-1 mice (13), which utilized the synthetic cannabinoid benchmark compound JWH-018. Additionally, earlier studies on the reinforcing effects of 5F-MDMB-PICA in CD-1 mice (22) guided our selection of a dosage range between 1 and 5 μg/kg/25 μl infusion.

The first key finding of our study highlights that 5F-MDMB-PICA has significant reinforcing properties in the IVSA paradigm used to determine drug reinforcement in adolescent CD1 mice, as demonstrated by their acquisition of operant behavior (lever pressing) with a higher number of active vs. inactive lever presses at the intermediate dose of 2.5 µg/kg/inf. (Fig. 2 A). The significant increase in drug intake observed in mice receiving the intermediate dose (2.5 μg/kg/inf., Fig. 2 B) compared to that observed with the doses of 1 and 5 μg/kg/inf., suggests a dose-dependent preference similar to the addiction profiles seen in other studies with the prototypical SCRA JWH-018 (13,21). Consistent with previous findings, the dose-response study showed an inverted U-shaped trend (13,21) (Fig. 2 A, B). Once the dose necessary and sufficient to acquire and sustain operant behavior was identified (2.5 μg/kg/inf.), we tested adolescent mice under different FR and PR schedules of reinforcement to better characterize the strength of the reinforcing properties of 5F-MDMB-PICA and highlight its abuse potential (PND 40-55). We observed that as the FR schedule of reinforcement was increased (from 1 to 3), the difference between active and inactive lever presses became greater (Fig. 2 C) while this effect was absent in the Vehicle group (Fig. 2 D). Importantly, operant behavior was specifically directed at obtaining 5F-MDMB-PICA, as shown by the greater number of active lever presses under PR schedules of reinforcement (Fig. 2 C).

**Fig. 1:**
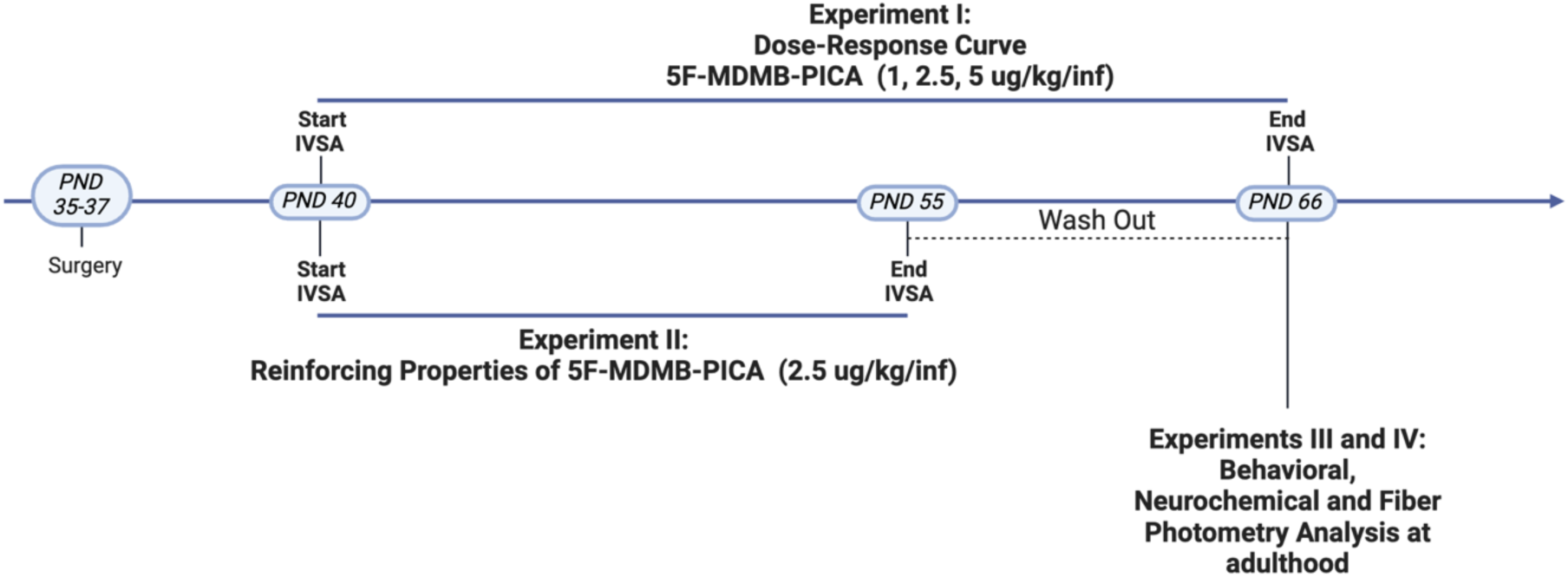
Experimental Timeline. The diagram illustrates the timeline of the experiments conducted on adolescent mice. Post Natal Day (PND), intravenous self-administration (IVSA).

**Fig. 2:**
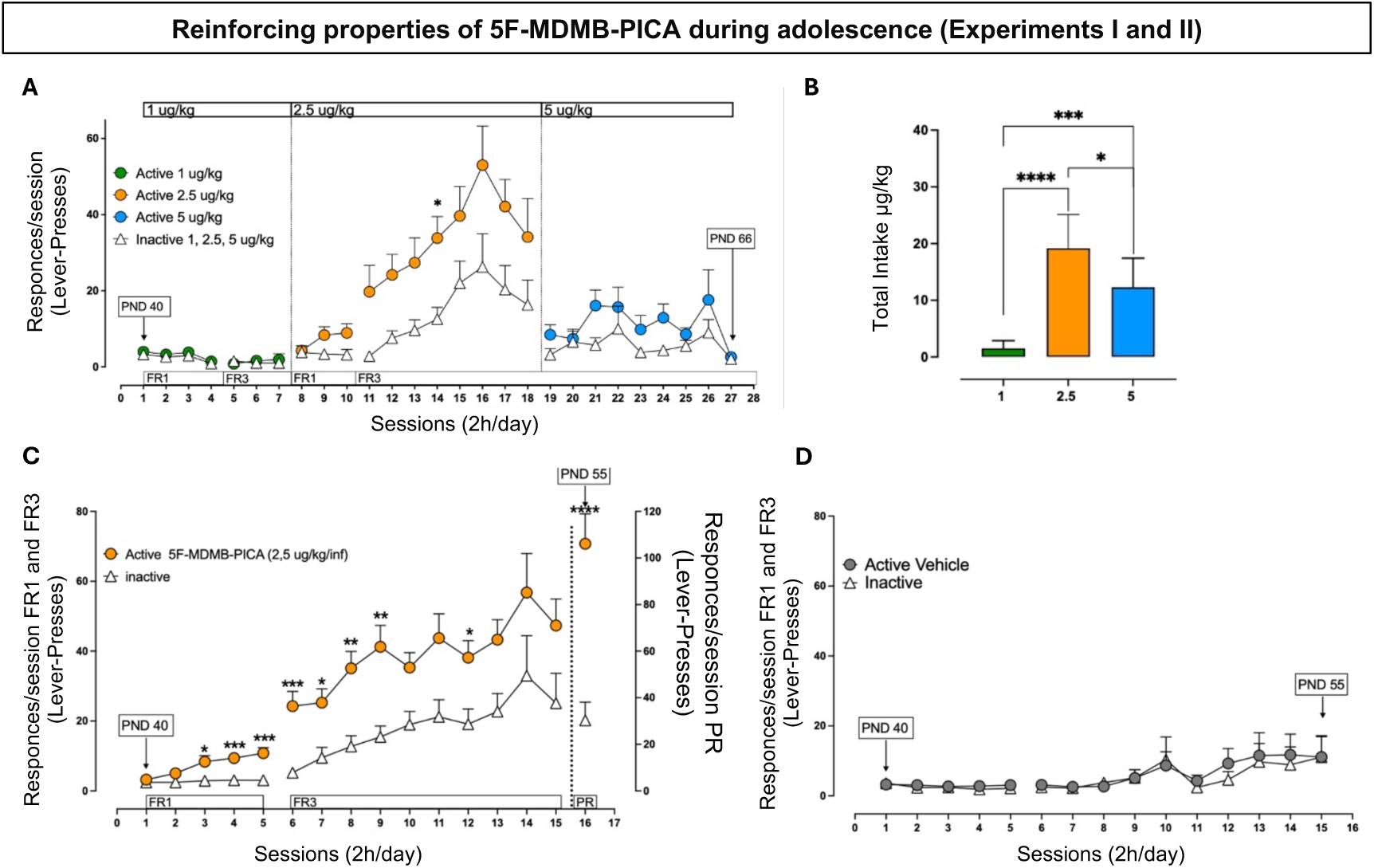
Reinforcing properties of 5F-MDMB-PICA during adolescence (Experiments I and II) **(A)** 5F-MDMB-PICA dose-response curve in the IVSA experimental paradigm. Results are expressed as mean ± SEM of the numbers of active (circles) and inactive (triangles) lever presses exhibited during each 2 h daily session under FR1 and FR3 reinforcement schedules at different 5F-MDMB-PICA doses 1 (active lever in green), 2.5 (active lever in orange), 5 (active lever in blue) μg/kg/inf. Two-way RM ANOVA or REML analysis (response × session) of the active vs inactive lever pressed performed at the dose of 1, 2.5 and 5 μg/kg/inf under FR-1 and FR-3 schedules revealed significant effects of session as follows: FR-1 at the dose of 1 μg/kg/inf (F(2.062, 41.23) = 5.368, p < 0.01, η² ≈ 0.212) (Two-way RM ANOVA); FR-3 at the dose of 5 μg/kg/inf (F(2.787, 53.65) = 3.002, p = 0.0418, η²p ≈ 0.135) (REML); FR-3 at the dose of 2.5 μg/kg/inf (F (4.068, 79.04) = 5.851”, p = 0,0003, η2p ≈ 0.232) (REML); and response as follows: FR-3 at the dose of 2.5 μg/kg/inf (F (1, 20) = 12.68, p = 0,0020, η2p ≈ 0.388) (REML). Sidak’s post hoc test of response and session main effect showed a significant difference between active and inactive lever presses at the dose of 2.5 μg/kg/inf in the 14th session (p < 0,05, t = 3.313, d ≈ 3.31); solid symbols: *p<0.05, vs inactive lever presses. N = 11. **(B)** Total intake during 5F-MDMB-PICA IVSA dose–response curve. Data are expressed as μg/kg of 5F-MDMB-PICA self-administered during each 2 h daily session. Each bar represents the mean ± SEM of the drug self-administered at each dose. Ordinary one-way ANOVA analysis revealed a significant effect of 5F-MDMB-PICA dose (F (2, 26) = 28.54, P<0.0001). Tukey’s multiple comparisons post hoc test of 5F-MDMB-PICA dose showed higher 5F-MDMB-PICA intake at the dose of 2.5 μg/kg/inf as compared to the doses of 1 (p < 0.0001), and 5.0 (p < 0.05) μg/kg/ inf. Solid symbols: *p<0.05, ***p<0.001, **** p< 0.0001 vs all the other groups. N= 11. **(C)** Response rates in different reinforcement IVSA schedules (FR1-FR3-PR). Results are expressed as mean ± SEM of the numbers of active (orange circles) and inactive (white triangles) lever presses exhibited during each 2h daily session under FR1, FR3 or PR reinforcement schedules during 5F-MDMB-PICA (2.5 µg/kg/inf) IVSA. Under FR1 schedule, two-way RM ANOVA analysis (response × session) revealed a significant main effect of session (F (2.742, 202.9) = 10.04, p < 0.0001, η2 = 0.049), response (F (1, 74) = 17.75, p < 0.0001, η2 = 0.108), and of response × session interaction (F (4, 296) = 6.803, p < 0.0001, η2 = 0.033) were observed. Sidak’s post hoc test revealed significant differences between active and inactive lever presses in the 3th (p < 0.05, t = 2,898, d ≈ 0.47), 4th (p < 0.001, t = 4,114, d ≈ 0.67) and 5th (p < 0.0001, t = 4,692, d ≈ 0.76) sessions. Under FR3 schedule, Mixed-effects model (REML) analysis (response × session) showed a main effect of response (F (1, 74) = 16.52, p = 0.0001, η² ≈ 0.53), and session (F (2.791, 191.1) = 8.095, p < 0.000, η² ≈ 0.41). Sidak’s post hoc test of response revealed significant differences between active and inactive lever presses in the 6th (p < 0.001, t = 4,340, d ≈ 0.70), 7th (p < 0.05, t = 3,185, d ≈ 0.52), 8th (p < 0.005, t = 3,895, d ≈ 0.63), 9th (p < 0.005, t = 3,733, d ≈ 0.61) and 12th (p < 0.05, t = 2,936, d ≈ 0.48) sessions. Unpaired t-test revealed significant differences between active and inactive lever presses during PR schedule (p < 0.0001, t = 5.033, df = 18). Solid symbols: *p<0.05, ** p< 0.01, *** p< 0.001, **** p< 0.0001 vs inactive lever presses; FR1 and FR3 N= 38, PR N= 10. **(D)** Number of responses on the active lever (circles) that resulted in Vehicle infusion (25 μl/inf) and on the inactive lever (triangles) during each 2h daily session under FR1, FR3 reinforcement schedules. Results are expressed as mean ± SEM; N= 25.

### Long-Term Behavioral Impact of Adolescent 5F-MDMB-PICA IVSA: Increased Aggression, Reduced Sociability, Anhedonia, and Increased Aversive Behaviors in Adult Mice

To evaluate the long-term behavioral effects of adolescent 5F-MDMB-PICA IVSA, our study employed three distinct behavioral tests: Resident Intruder Test (RIT), Sucrose Preference Test (SPT), and Olfactory Reactivity Test (ORT) (PND 66-78). These tests were chosen to assess various aspects of behavior, including social aggression, anhedonia, and responses to anxiety-inducing stimuli.

Given the extensive correlation between SCRA consumption and subsequent increases in aggressive behavior in both humans (23–27) and animal models (28,29) along with findings that correlate chronic SCRA treatment with persistent impairments in social behavior (30,31), we tested whether similar behavioral patterns could be observed in adulthood after adolescent IVSA of 5F-MDMB-PICA. To this end, we employed the RIT which is a well-established model for assessing territorial aggression and social interactions in rodents (32). RIT revealed that mice exposed to 5F-MDMB-PICA during adolescence exhibited higher aggression and fewer social interactions compared to Vehicle-treated controls, indicating increased aggressive behavior and reduced sociability (Fig. 3 A).

**Fig. 3:**
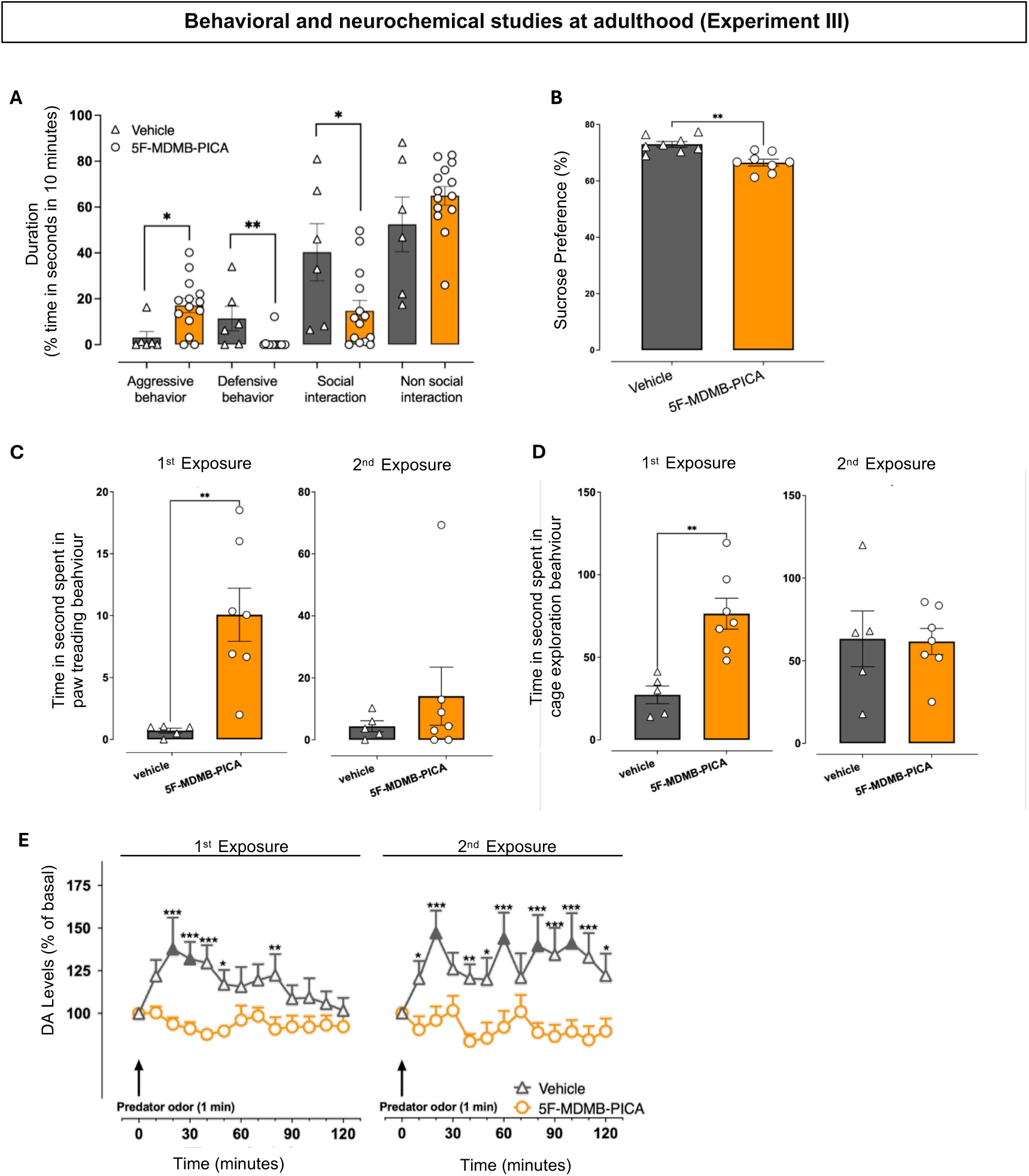
Behavioral and neurochemical studies at adulthood (Experiment III) **(A)** Aggressive, defensive behaviors and social and non-social interactions in response to a male intruder: Results are expressed as mean ± SEM of the % of time spent in each class of behaviors. Mann-Whitney U test revealed a significant increase in the time spent in aggressive behavior in the 5F-MDMB-PICA-treated group (U = 13, p < 0.05, r = 0.155) and a significant reduction in the time spent in defensive behavior (U = 12, p < 0.005, r = 0.143) as compared to the vehicle group. Additionally, two-tailed unpaired t-test revealed a significant reduction in the social interaction in the 5F-MDMB-PICA-treated group (95% CI: -47.51 to -3.55, p < 0.05, t = 2.440, df = 18, R² = 0.248) as compared to the vehicle-treated group. The analysis of non-social interaction scores showed no significant differences between the groups. Aggressive and defensive behaviors: solid symbols: *p<0.05; **p<0.005 Vehicles-treated group vs 5F-MDMB-PICA-treated group (Mann-Whitney test); Social interaction: solid symbols: *p<0.05 Vehicles-treated group vs 5F-MDMB-PICA-treated group (Unpaired t-test). N= 6 (Vehicle); 14 (5F-MDMB-PICA). **(B)** Anhedonic-like emotional state: The results of the sucrose preference test administered 10 days after the last IVSA session demonstrated a decreased sucrose preference in the 5F-MDMB-PICA-treated group as compared with the control mice. Results are expressed as mean ± SEM of the % of sucrose preference. An unpaired t-test was used to compare the sucrose preference scores between the vehicle-treated group and the 5F-MDMB-PICA-treated group. The analysis revealed a significant reduction in sucrose preference in the 5F-MDMB-PICA-treated group (66.49%) compared to the vehicle-treated group (72.97%), with a mean difference of -6.482% (95% CI: -9.894 to -3.070, p < 0.005, t = 4.075, df = 14, R² = 0.5426). Solid symbols: **p<0.005 5F-MDMB-PICA-treated group vs Vehicles-treated group; N = 8 (Vehicles-treated group); 8 (5F-MDMB-PICA-treated group). **(C) (D)** Effects of the odor predator exposure (2-Methyl-2-propanethiol inhalation) on Paw Treading (C) and cage exploration (D) behaviors in adult mice that underwent intravenous self-administration of 5F-MDMB-PICA or Vehicle during adolescence: Results are expressed as mean ± SEM of the time spent in each behavior. Unpaired t-test revealed a significant increase in paw trading behavior (A) and cage exploration behavior (B) during the 1st exposure in the 5F-MDMB-PICA-treated group as compared to the vehicle-treated group (paw trading behavior: 95% CI: 3.609 to 15.11, p < 0.005, t = 3.626, df = 10, R² = 0.568); (cage exploration behavior: 95% CI: 22.19 to 76.04, p < 0.005, t = 4.065, df = 10, R² = 0.6229). The analysis of the 2nd exposure didn’t show any significance for both the behaviors. **p<0.005 Vehicle vs 5F-MDMB-PICA-treated group; N= 5 (Vehicle-treated group 1st and 2nd exposure); 7 (5F-MDMB-PICA-treated group 1st and 2nd exposure). **(E)** Adolescent 5F-MDMB-PICA IVSA induces adaptive changes in mPFC dopamine responsiveness to repeated exposure to a stressful odor stimulus (2-Methyl-2-propanethiol inhalation): data are presented as the mean ± SEM of changes in extracellular dopamine in the mPFC, expressed as the percentage of basal values. The arrows indicate the start of the 1st and 2nd odor exposure (1 minute). Two-way RM ANOVA analysis of the <u>1st odor exposure</u> revealed a significant effect of treatment (F (1, 16) = 9.743, p < 0.01, η2 ≈ 0.183) and time × treatment interaction (F (12, 192) = 2.089, p < 0.05, η2 ≈ 0.055). Tukey’s multiple comparisons post hoc test revealed an increase of dialysate dopamine in the mPFC of the Vehicle-treated group as compared to basal values at minutes 20 (p< 0.005) and 30 (p < 0.05) and as compared to 5F-MDMB-PICA-treated group at minutes 20 (p < 0.0005), 30 (p < 0.0005), 40 (p < 0.0005), 50 (p < 0.05), 80 (p < 0.01). Two-way ANOVA analysis of the <u>2nd odor exposure</u> revealed a significant effect of treatment (F (1, 13) = 14.08, p < 0.005, η2 ≈ 0.280) and a time × treatment interaction (F (12, 156) = 2.244, p < 0.05, η2 ≈ 0.049). Tukey’s multiple comparisons post hoc test revealed an increase in dialysate DA in the mPFC of the vehicle group as compared to basal values at minutes 20 (p < 0.005), 60 (p < 0.05), 80 (p < 0.05), 100 (p < 0.05) and as compared to 5F-MDMB-PICA-treated group at minutes 10 (p < 0.05), 20 (p < 0.0005), 40 (p < 0.01), 50 (p < 0.05), 60 (p < 0.0005), 80 (p < 0.0005), 90 (p < 0.001), 100 (p < 0.0005), 110 (p < 0.001), 120 (p < 0.05). Solid symbols: full triangles or circles p<0.05 or 0.005 vs the basal values; * p<0.05, ** p<0.005, *** p< 0.0005 vs Vehicles. N= 9 (Vehicle-treated group 1st exposure); 9 (5F-MDMB-PICA-treated group 1st exposure); 6 (Vehicle-treated group 2nd exposure); 9 (5F-MDMB-PICA-treated group 2nd exposure).

Moreover, given the well-documented long-term emotional deficits associated with the administration of SCRAs during development (33–35), we investigated how adolescent IVSA of 5F-MDMB-PICA affects the ability to experience pleasure in adulthood. To do this we used the SPT, a reward-based test used as a measure of anhedonia (36). The SPT assesses an animal’s preference for a sweet solution over water. Our findings showed a decreased preference for a 1% sucrose solution in mice that self-administered 5F-MDMB-PICA during adolescence (Fig. 3 B), suggesting that these animals have a compromised ability to experience pleasure, indicative of an anhedonic emotional state in adulthood.

Finally, when assayed for ORT, adolescence mice that self-administered 5F-MDMB-PICA, demonstrated significant behavioral changes indicating a persistent aversive emotional state. The ORT was conducted to assess the behavioral responses of Vehicle and 5F-MDMB-PICA mice to two consecutive exposures to an aversive odorant (i.e. 2-methyl-2-propanethiol). At the 1^st^ exposure, 5F-MDMB-PICA mice exhibited significantly higher paw treading behavior scores (Fig. 3 C), a well-known indicator of aversion in rodents (37,38), and spent significantly more time engaging in cage exploration behaviors compared to the Vehicle group (Fig. 3 D). No significant differences in these behaviors between groups were observed during the 2^nd^ exposure (Fig. 3 C, D). This observation aligns with the trend noted during the RIT conducted in this study, where 5F-MDMB-PICA self-administered mice displayed a propensity for more non-social interactions than the Vehicle group.

These results collectively demonstrate that adolescent exposure to 5F-MDMB-PICA leads to long-term behavioral changes in adulthood, including increased aggression, reduced sociability, anhedonia, aversive emotional state and related behaviors.

### Adolescent IVSA of 5F-MDMB-PICA Leads to Dysregulation of DAergic System and Neuronal Excitability in the mPFC in Response to Olfactory Aversive Stimuli in Adulthood

Given the behavioral changes observed, we examined the neurochemical and neurophysiological mechanisms underlying these behaviors in adult mice previously exposed to 5F-MDMB-PICA during adolescence. It is well known that the mPFC plays a significant role in behavioral control and addiction (39–45). Therefore, we decided to investigate the functionality of this critical region by employing microdialysis and fiber photometry during the Olfactory Reactivity Test to provide a deeper understanding of the long-term neurobiological consequences of early 5F-MDMB-PICA exposure.

Our microdialysis results during the ORT underscore significant alterations in DA responsiveness in the mPFC (Fig. 3 E). Indeed, mice exposed to 5F-MDMB-PICA during adolescence showed blunted DA responses during repeated exposures to the aversive olfactory stimulus compared to the Vehicle group, which exhibited typical increases in DA levels, indicative of a normal response to a stressor (Fig. 3 E).

Complementing our neurochemical findings, we employed fiber photometry recording using the calcium sensor GCaMP, which provided insights into the real-time neuronal activity of excitatory neurons within the mPFC during exposure to the same aversive stimulus. Control animals displayed a sharp increase in calcium activity during the first and second exposure, which is consistent with the expected activation of the mPFC in response to stress (Fig. 4). In contrast, mice that previously self-administered 5F-MDMB-PICA showed reduced neuronal activity in the 1^st^ but not in the 2^nd^ odor exposure (Fig. 4). This diminished responsiveness not only supports our microdialysis results but also suggests a broader alteration in the functional output of mPFC neurons. These diminished neural responses may reflect a disruption in the neural circuitry involved in emotional processing and stress regulation, potentially contributing to the behavioral phenotypes observed, such as increased aggression, reduced social interaction, and an anhedonic state. This hypothesis is supported by research indicating that alterations in mPFC activity can significantly impact behavioral and emotional regulation (20,46,47).

**Fig 4:**
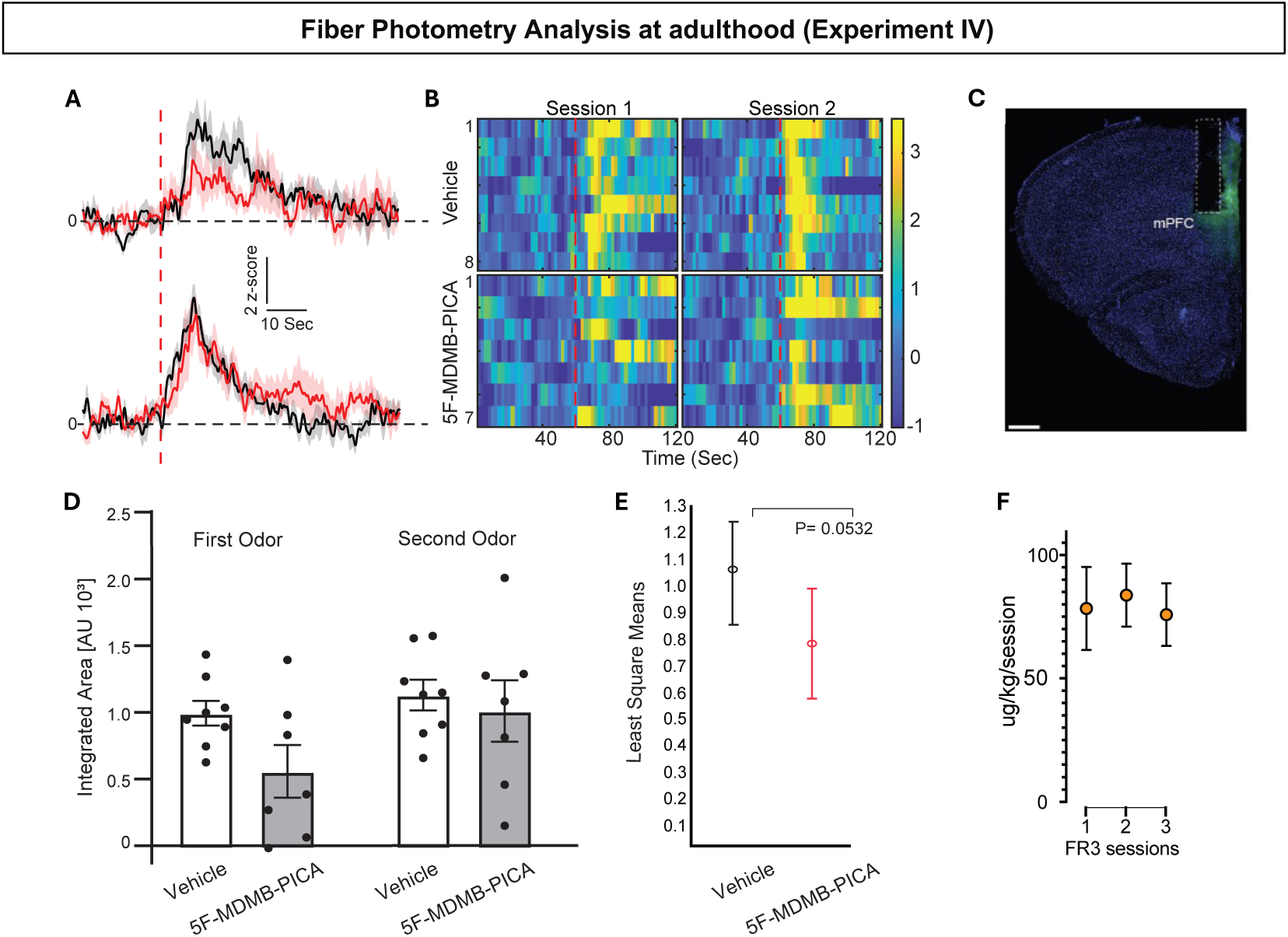
fiber Photometry Analysis at adulthood (Experiment IV) **(A)** Average trace of mice after self-administration of 5F-MDMB-PICA (red) or vehicle (black) during the 1st odor exposure (top panel) and 2nd odor exposure (bottom panel). The red dotted line indicates when mice were exposed to the odorant. Scale bars demonstrate 2 z-scores from the mean (y-axis) and 10 seconds in duration (x-axis). Shaded regions indicate ± SEM. **(B)** Heatmap of z-scored calcium activity across all vehicle mice (y-axis, top rows) and 5F-MDMB-PICA mice (bottom rows) over time, the red dotted line represents odorant exposure. The left panel displays the 1st exposure while the right demonstrates the 2nd. Z-scored changes in fluorescent activity range from -1 to 3. **(C)** A representative image displaying the location of the optic fiber cannulae and GCaMP expression. **(D)** Integrated area of calcium traces in arbitrary units (AU) of 15 seconds following odor presentations (range from 5 seconds after odor presentation to 20 seconds after) per mouse during the first and second odor exposure. Bars represent the mean ± SEM. **(E)** Least squares means from the linear mixed model (LMM), which refers to the predicted means for each treatment group, adjusted for other terms in the model, comparing odor-induced calcium activity in vehicle-treated mice versus 5F-MDMB-PICA-treated mice. 5F-MDMB-PICA IVSA treatment resulted in an approximately 26.14% reduction in neuronal activity compared to the vehicle treatment (treatment effect, F(1,13) = 4.5, P = 0.0532). We did not detect a significant effect of exposure time (F(1,13) = 2.4, P = 0.14) or an interaction between exposure time and treatment groups (F(1,13) = 0.69, P = 0.418). **(F)** Data summarizing the amount of 5F-MDMB-PICA self-administered during the 3 FR3 sessions (mean ± SEM).

## Discussion

Our study demonstrates for the first time that adolescent exposure to 5F-MDMB-PICA results in persistent neurobehavioral alterations, likely mediated by disruptions in DAergic signaling and mPFC excitability, as evidenced by: i) altered behavioral outcomes in the resident intruder test, indicative of increased aggression and antisocial behavior; ii) the induction of an anhedonic emotional state, as indicated by the sucrose preference test; iii) increased aversive behavioral responses to socially intimidating stimuli, such as predator odor, in the odor reactivity test; and iv) alterations in neuronal pathways in the mPFC, involving changes in DA levels and neuronal excitability within the mPFC. These findings emphasize the heightened vulnerability of the adolescent brain to SCRAs and highlight the urgent need for further research into their long-term consequences.

### Reinforcing Properties of 5F-MDMB-PICA during adolescence

Our study demonstrates that 5F-MDMB-PICA exhibits reinforcing effects in adolescent mice, as evidenced by their acquisition of operant behavior in the IVSA paradigm. Notably, the intermediate dose (2.5 μg/kg/inf) was the most effective in maintaining self-administration, consistent with the inverted U-shaped dose-response patterns observed for other synthetic cannabinoids, including JWH-018 (13,21,48). Interestingly, this effective dose was nearly three times lower than that required for JWH-018, potentially reflecting the higher affinity of 5F-MDMB-PICA for CB1 and CB2 receptors (17,49). The progressive ratio paradigm further confirmed the motivational salience of 5F-MDMB-PICA, with adolescent mice exhibiting persistent drug-seeking behavior even under increased response demands. These findings underscore the abuse liability of 5F-MDMB-PICA and suggest that its potency may contribute to compulsive drug-taking behaviors observed in human SCRA users (5,50).

### Long-Term Behavioral Impact of Adolescent 5F-MDMB-PICA IVSA: Increased Aggression, Reduced Sociability, Anhedonia, and Increased Aversive Behaviors in Adult Mice

Beyond its reinforcing effects, adolescent exposure to 5F-MDMB-PICA induced persistent behavioral changes in adulthood. Our findings in the resident-intruder test revealed a significant increase in aggressive behaviors and a reduction in social interactions in adult mice exposed to 5F-MDMB-PICA during adolescence, aligning with reports linking SCRA consumption to heightened aggression in humans (23–25) and animal models (28,29). Case studies have documented aggression following Spice/K2 consumption (23), while epidemiological studies indicate increased violent behaviors among adolescents using SCRAs compared to THC users (24). While extensive literature underscores the correlation between chronic cannabis consumption and aggressive behavior in humans (25–27), few studies have explored this relationship in animal models. Preclinical studies focusing on acute cannabis or SCRA exposure report conflicting results (51–55). Moreover, it has been suggested that the loss of CBR function leads to increased aggression, suggesting that endocannabinoid (eCB) signaling may play an inhibitory role in aggressive tendencies (56–58). Our findings indicate that 5F-MDMB-PICA exposure during adolescence disrupts emotional and motivational circuits, potentially through alterations in the eCB system, which plays a crucial role in brain development and social behavior regulation (59,60). The eCB system modulates neurotransmitter release and synaptic plasticity (61); thus, its disruption during critical developmental stages can lead to long-lasting changes in brain function and behavior.

In addition to aggression, the RIT allowed us to observe reduced sociability in adult mice that self-administered 5F-MDMB-PICA in adolescence, consistent with previous findings on chronic SCRA exposure (30). Chronic treatment with WIN 55,212-2 during adolescence has been linked to persistent social impairments resembling schizophrenia symptomatology (33–35) and social deficits persisted even after treatment cessation with the antipsychotic quetiapine. Other studies have similarly reported SCRA-induced sociability impairments (31), reinforcing the evidence of long-term detrimental effects of SCRAs on social behavior.

Moreover, mice that self-administered 5F-MDMB-PICA exhibited a persistent anhedonic state in adulthood, as reflected by reduced sucrose preference in the sucrose preference test. This aligns with reports linking chronic SCRA exposure to anhedonia and emotional blunting (34). Our results add to this body of evidence, suggesting that adolescent SCRA consumption disrupts not only aggressive and social behaviors but also emotional processing.

Altered responses to aversive odorant stimuli further support the presence of an emotional dysregulation phenotype. Mice previously exposed to 5F-MDMB-PICA exhibited heightened stress responses, as evidenced by increased paw treading and cage exploration during the first exposure to an aversive odor. These findings align with observations in JWH-018-treated rats, where taste neophobia and an aversive state were noted during the first but not the second exposure to chocolate (62).

Our comprehensive behavioral analysis revealed increased aggression, reduced social interactions, anhedonic state, and a heightened aversion to aversive odorant stimuli in mice exposed to 5F-MDMB-PICA during adolescence. These alterations, evidenced by the RIT, SPT, and ORT, likely stem from SCRA-induced dysregulation of the eCB system, which plays a pivotal role in anxiety and stress modulation (63,64).

### Adolescent IVSA of 5F-MDMB-PICA Leads to Dysregulation of DAergic System and Neuronal Excitability in the mPFC in Response to Olfactory Aversive Stimuli in Adulthood

To understand the neurobiological basis of these behavioral alterations, we examined DA release and neuronal excitability in the mPFC of adult mice previously exposed to 5F-MDMB-PICA. The mPFC is crucial for emotional and stress-related responses, with dysfunction implicated in several neuropsychiatric disorders (40–45).

Our microdialysis results indicate impaired stress-related DA signaling in 5F-MDMB-PICA-exposed mice, as shown by blunted DA responses, in the mPFC, to aversive odorant exposure as compared to the typical stress-induced DA increases observed in controls. This aligns with studies indicating that chronic SCRA exposure during critical developmental periods dysregulates DAergic function (62,65) and can lead to lasting changes in the neurochemical systems that regulate emotion and stress responses, even if research into the effects of repeated SCRA exposure on DA activity in the mPFC is limited.

Notably, repeated JWH-018 administration has been shown to decrease VTA DA neuronal activity, with persistent effects post-withdrawal (62), highlighting a profound dysregulation of the DA system. Chronic SCRAs exposure has been linked to reduced DA responses in the mPFC and NAc to both drug and non-drug stimuli, likely contributing to the dysregulation of motivation and emotional processing (62). Moreover, several researchers have highlighted the critical role of DAergic projections from the VTA to the mPFC in response to stress or aversive cues (66–70). DA habituation to repeated non-drug stimuli, such as chocolate, was also observed (65), an effect previously linked to impaired top-down control from the mPFC over subcortical DAergic areas (71). These findings confirm the central role of the mPFC in behavioral regulation and suggest a lasting impact of adolescent SCRA exposure on DA system function.

Additionally, SCRAs have been associated with CB1R downregulation in DA-related areas, supporting the hypothesis that CB1R desensitization occurs following prolonged SCRA exposure (56,72,73). These observations support the hypothesis that eCB signaling plays a critical role in the DA dysregulation we observed, underscoring the significant impact of sustained SCRAs exposure on neural pathways involved in reward and motivation.

Complementary fiber photometry results revealed diminished neuronal excitability in the mPFC during the first aversive odorant exposure in 5F-MDMB-PICA IVSA mice, reinforcing the notion that adolescent SCRA exposure disrupts stress-processing mechanisms.

Indeed, control animals displayed a sharp increase in calcium activity during the first and second odorant exposure, which is consistent with the expected stress response of the mPFC. In contrast, mice that self-administered 5F-MDMB-PICA showed significantly reduced neuronal activity during the first exposure. This diminished responsiveness not only supports our microdialysis results but also suggests a profound alteration in the functional output of mPFC neurons. These diminished neural responses may reflect a broader disruption in the neural circuitry involved in emotional processing and stress regulation, potentially contributing to the behavioral phenotypes observed, such as increased aggression, reduced social interaction, and increased anhedonia.

These effects may be linked to altered receptor sensitivity or changes in DAergic neuron function. Moreover, the infra limbic (IL) region of the mPFC plays a key role in suppressing aversive behaviors via inhibition of amygdalar output (74,75). Reduced activity in this area has been linked to deficits in cognitive control and emotional regulation (76–78), highlighting the complexity and significance of mPFC function in emotional regulation and control of behaviors. Nevertheless, VTA DAergic projections to specific neurons within mPFC can be distinguished by their location in the dorsomedial VTA and their specific electrophysiological and functional responses to aversive stimuli, as well as their lack of D2 autoreceptors, emphasizing their unique role in DAergic neurotransmission (79).

The discrepancy between microdialysis and fiber photometry findings regardless of the divergent response observed during the second exposure may stem from differences in temporal resolution and neurochemical specificity. While microdialysis revealed no significant increases in DA levels in the mPFC of 5F-MDMB-PICA-exposed mice during both odor exposures, fiber photometry showed that mPFC excitatory neurons exhibited a reduced response during the first exposure but showed no difference in activation compared to controls during the second exposure. This delayed response may reflect neuronal adaptation or synaptic plasticity mechanisms underlying stress processing (80–82).

Taken together these findings suggest that adolescent exposure to 5F-MDMB-PICA induces persistent DAergic and neuronal dysregulation within the mPFC, potentially contributing to the observed behavioral alterations. The blunted DA responses and altered neuronal activity indicate a possible desensitization of the mPFC to stress-related stimuli, consistent with broader evidence implicating eCB signaling in DA dysregulation following chronic CB exposure. These neuroadaptive changes in the mPFC, including modifications in DA receptor expression, synaptic plasticity, and neuronal excitability, likely play a crucial role in the long-term behavioral effects of SCRAs. Overall, our findings reinforce the notion that adolescent SCRA exposure can induce lasting disruptions in brain function, with significant implications for emotional and cognitive health. Understanding the mechanisms underlying SCRA-induced behavioral dysregulation is critical for developing effective strategies to prevent and treat the adverse effects associated with adolescent substance use.

## Supporting information

Supplementary informations

## Acknowledgement

This research has been funded by MUR with “Progetti di Rilevante Interesse Nazionale (PRIN-PNRR) 2022”, project: “DECODE-018-Dissecting the enduring changes in the prefrontal cortex induced by exposure to the synthetic cannabinoid JWH-018 during adolescence: multidisciplinary characterization of the behavioral, neurochemical, and molecular outcomes at adulthood in rats and mutant mice (P20229TKXR; PI: Prof. MA De Luca); the Drug Policies Department, Presidency of the Council of Ministers, Italy, project: “Implementation of the detection and study of the effects of NPS: Development of a multicentric research team for the enhancement of the National Observatory of Dependence and of the National Early Warning System” (CUP: I55E22000320001; Unit Leader for University of Cagliari, Prof. MA De Luca).

