## Supplementary informations for "Long-Term Effects of Adolescent 5F-MDMB-PICA Intravenous self-administration: Neurobehavioral Consequences and medial Prefrontal Cortex Dysfunction in Adult Mice"

### Materials and methods

#### Drugs and chemicals

5F-MDMB-PICA was purchased from Cayman Chemical Company (Ann Arbor, MI, USA) and dissolved in 0.5% EtOH, 0.5% Tween 80 and 99% saline.

The molecule chosen as an aversive olfactory stimulus, 2-methyl-2-propanethiol, is a thiol consisting of a methyl group (-CH3) and a thiol group (-SH) bound to the same tertiary carbon atom. This compound shares a structural similarity with 3-methyl-1-butanethiol, previously demonstrated to be significantly aversive to mice and found in the urine of female and male red fox (1,2). The rationale for using this substance is to assess the response of the mice mPFC toward sulfur-containing odorants known to be similar to components in the odor of natural predators of the mouse. The 2-methyl-2-propanethiol was obtained from Sigma–Aldrich (CAS Number: 75-66-1), ensuring a nominal purity of at least 99% and was diluted 1:10000 in distilled $H_2O$.

#### (i) IVSA studies during adolescence (Experiments I and II)

##### Surgical Procedures for Catheter Placement

On PND 35-37, mice were anesthetized with isoflurane gas, and maintained under anesthesia using a breathing tube under a scavenging system. Lubricant eye drops (Tears Naturale, Alcon, USA) were applied to both eyes to prevent them from drying out during the procedure. A chronic intravenous catheter for mice was prepared and implanted in the right jugular, as previously described (3,4). After surgery, mice were housed individually under standard conditions; in order to ensure full recovery, mice underwent postoperative antibiotic treatment with gentamicin sulphate (30 mg/kg, subcutaneously [s.c.]), for 3 days to aid recovery before starting the IVSA.

**Intravenous Self-Administration procedures (Experiments I and II)**

Daily two-hour SA sessions were performed 7 days/week for: 27 days, PND 40-66 (Experiment I); 16 days, PND 40-55 (Experiment II); and 15 days, PND 40-54 (Experiment IV) in Skinner boxes equipped with soundproof insulation (model: LE 26M, Panlab Harvard Apparatus, Spain for experiment I and II; and Med Associates inc apparatus, Fairfax, Vermont, USA for experiment IV). Within the chamber, the operant panel was equipped with two levers, one active (right) and one inactive (left). The active lever was associated with the administration of 25 µL of 5F-MDMB-PICA (1-5 µL/kg), while the inactive lever had no associated consequences. A white light served as discriminative stimuli over the active lever. Prior to each daily SA session, the mice's catheter was flushed with 0.1 ml of saline solution, and post-session, it was flushed with 0.1 ml of heparinized saline (50U/ml). Additionally, a 4 g food pellet was provided after each daily SA session to maintain stable body weights. The mice performance in each session was recorded using Panlab software (Panlab Harvard Apparatus, Spain for experiments I and II and MedPC software, Fairfax, Vermont, USA for experiment IV). The experimental protocol encompassed distinct phases designed to regulate the administration of the substance following the setup previously described by (5).

**Characterization of 5F-MDMB-PICA dose-response curve in the IVSA experimental paradigm (Experiment I)**: To identify the dose-response curve of 5F-MDMB-PICA we used a range of doses from 1 to 5 (μg/kg/25 ul inf.). From PND 40 to PND 43 and from PND 47 to 49, sessions were carried out under FR1 schedule of reinforcement (1 active lever press: 1 injection) at the doses of 1 and 2.5 (μg/kg/25 ul inf.) respectively. From PND 44 to 46 and from PND 50 to 66 the protocol was switched to FR3 schedule of reinforcement (3 active lever press: 1 injection). The criterion for transitioning from FR1 to FR3 was based on the methodology described by (5). Sessions were performed using increasing doses of 5F-MDMB-PICA from the lower 1 (μg/kg/25 ul inf.), intermediate 2.5 (μg/kg/25 ul inf.) till the higher 5 (μg/kg/25 ul inf.). During the sessions at each infusion followed a 20-sec time out period, in this time no more drug was made available and the light above the active lever was shut down, however in the meantime all the activity on the levers were still recorded.

**Characterization of Response rates in different reinforcement IVSA schedules (FR1, FR3, PR) (Experiment II)**: Identified the dose (2.5 μg/kg/25 ul inf.) at which mice acquired the operant behavior we used different FR protocols (5 sessions FR1;10 sessions FR3; 1 session Progressive Ratio (PR)) to evaluate the abuse properties of 5F-MDMB-PICA. The duration of the protocol was designed based on similar studies previously conducted in our laboratory using JWH-018 as a reference compound of the synthetic cannabinoids, (Margiani at al., 2022). In the PR session each subsequent injection was based on the adapted exponential sequence: 1, 2, 4, 6, 9, 12, 15… (6) and the session lasted for 2 h.

The break point (BP) was defined as the highest ratio completed prior to the last injection. The same protocol was used for the vehicle group which performed IVSA with vehicle solution (2% ethanol, 2% Tween 20 and 96% saline) instead of 5F-MDMB-PICA solution.

**(ii) Behavioral and neurochemical studies at adulthood (Experiment III)**

**Resident intruder test**

Ten days after the drug-free period ended (PND 66), we employed the resident-intruder paradigm (7) to evaluate whether adolescent exposure to 5F-MDMB-PICA (2.5 μg/kg/25 μl infusion) was correlated with an increased propensity for aggressive behavior in adulthood. The test occurred in the resident's home cage (40 x 30 x 12 cm equipped with bedding material) in a sound attenuated room with the same light, temperature, and humidity of the house room. Behavior was videotaped and no other mice were present in the room during testing except for the resident and the intruder. The intruders were mice of the same strain, age, and housing conditions as the resident mice, but they had not undergone the IVSA sessions. After 30 minutes of acclimatization to the experimental room for the resident, the intruder was introduced into the room and the test was performed, the interactions between resident and intruder lasted 10 minutes. Videotapes capturing resident behavior were subsequently evaluated by an observer using a set of predefined behavioral parameters, as previously outlined (7).

The analyzed behaviors were categorized into four groups as described below:

- Offense behavior score: Sum of lateral threat, upright posture, clinch attack, keep down, and chase.

- Defense behavior score: Sum of flight, defensive upright posture, submission, and freeze.
- Social interaction score: Sum of social exploration, ano-genital sniffing, and social groom.
- Non-social exploration score: Sum of rest or inactivity, and cage exploration.

The recorded behavioral parameters accounted for 100% of the observation time, and the final score for each behavior was expressed as a percentage of the total observation time (7).

**Sucrose preference test**

The Sucrose Preference Test, as previously described (8), serves as a reward-based test to measure anhedonia, defined as the reduced capacity to feel pleasure. It utilizes the natural affinity rodents have for sweet flavors to assess any decline in attraction to a sucrose solution, which can be interpreted as anhedonia. For the test, at PND 66, mice were single housed in their regular cages provided with two different bottles: one containing plain water and the other a 1% sucrose solution over a period of 6 days, which includes 3 days for acclimatization (in which both bottles contain water) and 3 days of actual testing (in which in one bottle the water were replaced with sucrose solution). The two bottles' positions were switched every day, and the consumption of each liquid were recorded daily to determine the preference for sucrose. Preference for sucrose was determined by the ratio of sucrose solution consumed to the overall fluid consumption, expressed as a percentage, and averaged across the three days designated for testing. The formula for sucrose preference is:

$$\text{Sucrose Preference} = \frac{\text{total sucrose solution consumption}}{(\text{total water+total sucrose solution consumption})} \times 100$$

**Behavioral assay during Odor reactivity Test**

To minimize the number of animals used, the group of mice that underwent the sucrose preference test (PND 66–71) were allowed a recovery period of one week. Subsequently, on PND 78, these mice were subjected to behavioral assessment during the odor reactivity test.

The behavioral assay was conducted over two consecutive days and consisted of two phases: habituation, lasting 30 minutes, during which the mice were placed in an experimental cage (40 x 30 x 12 cm equipped with bedding material) with an odorless cotton stick; the test phase, lasting 6 minutes, involved presenting the same stick soaked with the odor solution (2-methyl-2-propanethiol) to the mice for 1 minute, followed by replacement with a cotton stick without odor for the remaining 5 minutes. The behaviors analyzed included interaction with the odor-stick (exploration latency, sniffing, biting, burying), aversive behaviors (paw treading, freezing), rearing, cage exploration, licking, and grooming. The duration of each behavior, during the 6 minutes test, was expressed in seconds.

**Neurochemical assay during Odor reactivity Test**

**In vivo brain microdialysis experiment**

In vivo brain microdialysis, a technique enabling the collection of samples from an animal's brain extracellular space while it moves freely, relies on substance diffusion between the brain's extracellular space and artificial cerebrospinal fluid

(ACSF - Ringer's solution) through a semipermeable membrane in a vertical microdialysis probe, equipped with an active dialyzing portion. This method, as described by Hernandez and colleagues (9) and Di Chiara (10), was employed in the present study to measure extracellular dopamine (DA) levels in the mPFC in response to an aversive stimulus in adult mice (PND 66) exposed to 5F-MDMB-PICA during adolescence, compared to vehicle-treated controls.

**Preparation of microdialysis probes**

Vertical microdialysis probes, with an active dialysing portion of 2 mm for mPFC were prepared with AN69 fibers (Hospal Dasco, Italy) as previously described (4).

**Surgical procedures for microdialysis probes placement**

Mice were anesthetized with isoflurane and then placed in a stereotaxic frame (Kopf Instruments, Tujunga, CA, USA) for the probe implantation. The probe was implanted into the medial PFC (mPFC), according to the Paxinos and Franklin mouse brain atlas (AP: ± 1.9; ML: ± 0.1; DV: -3.0 from Bregma) as previously reported (4,11).

**DA assessment during odor reactivity test**

One day after surgery, probes were perfused with Ringer's solution (composition in mM: 147 NaCl, 4 KCl, 2.2 CaCl2) at a constant rate of 1 ul/min. Dialysate samples (10 ul) were injected into an HPLC equipped with a reverse phase column (C8 3.5 microm, Waters, USA) and a coulometric detector (ESA, Coulochem II) to quantify dopamine (DA). The detector settings involved an oxidation electrode at +125 mV and a reduction electrode at -175 mV. The mobile phase comprised 50 mM NaH2PO4, 0.1 mM Na2-EDTA, 0.5 mM n-octyl sodium sulfate, 15% methanol (v/v), with a pH of 5.5, and the DA assay's sensitivity was determined to be 5 fmol/sample. After 2 h washing, basal DA levels were evaluated, and estimated as the mean of three consecutive samples whose values do not differ more than 10%. Then, mice were exposed to the odorous solution for 1 minute, by using a cotton stick soaked with the odor solution. The monitoring of extracellular DA levels continued for up to two hours, leading up to a second exposure to the odor. Following this second exposure, sample collection continued for an additional 2 hours.

**Hystology**

Upon completion of the microdialysis experiments, the animals were euthanized, and their brains were extracted and preserved in 8% formalin for histological analysis, ensuring the accurate placement of the microdialysis probes.

**(iii) Fiber Photometry Analysis at adulthood (Experiment IV)**

Fiber photometry is a cutting-edge technique that has revolutionized our ability to observe and understand the intricate dynamics of neuronal populations in living organisms. This approach leverages the specificity of genetically encoded calcium indicators (GECIs) to monitor the calcium dynamics within neurons, serving as a proxy for neuronal activity. Calcium ions play a pivotal role in neuronal signaling, and their concentration changes are indicative of neural activation or inhibition. By incorporating light to stimulate neurons and measuring the resultant fluorescence signals emitted by GECIs, fiber photometry provides a real-time, in vivo window into the functional activities and patterns of neural circuits. In the context of our study, we employed fiber photometry to investigate the effects of adolescent exposure to 5F-MDMB-PICA IVSA on the excitatory neurons activity within the mPFC in response to aversive stimuli. The experiment was carried out in collaboration with the team of Professor Francesca Telese at the University of California San Diego. CD-1 adolescent male mice (n=15), PND 30, were implanted with a jugular catheter for the administration of 5F-MDMB-PICA (by using the same method described previously) and in the same surgical session were also implanted with the optical fibers directly above the mPFC (AP: +1.75; ML: +0.4) via stereotaxic surgery. Additionally, to facilitate the monitoring of calcium dynamics within the mPFC, we injected a highly sensitive GCaMP calcium sensor (AAV-CamKIIa-jGCaMP8m, Addgene# 176751) into the same region. After 5 days of recovery, from PND 35 to PND 50 mice underwent the same IVSA protocol described above (5 days under FR1 and 10 days under FR3 schedule of reinforcement). Following a wash out period of 10 days neuronal activity was assessed during the odor reactivity test.

**Surgical Procedure for Catheter and fiber Implantation, and virus injections**

Mice were anesthetized with 5% isoflurane and then positioned in a stereotactic frame (RWD) and kept anesthetized by maintaining 1.5-2% isoflurane. The skull was exposed, and a small craniotomy was drilled over the mPFC implant site (AP: +1.75 ML: +0.4). A pulled glass pipette was loaded with a viral prep (AAV1-CaMKII-jGCAMP8m, Addgene 176751, final titer ~5x10E12) and it was injected using a nanoliter injector (Drummond Scientific, Nanoject III) into the mPFC (AP: +1.75 ML: +0.4 DV:-2.2 and -1.8, 200nl) in each site at a rate of 3nl/sec). Following injection, an optic fiber (RWD, NA 0.39, core 400um, Ferrule diameter 1.25) was positioned at 200-150 um dorsal to the lower injection site. Then, the skull was covered by a layer of Metabond C&B (Parkell), and a cap of dental cement (Ortho-

Jet, Lang dental) was built around the optic fiber ferrule). Mice were then returned to their home cages. Recording started at least 3 weeks following surgery to allow the virus expression.

#### Neuronal activity recording during odor reactivity test

On the day of the experiment, we used a commercial photometry system (Neurophotometrics, FP3002) to perform in-vivo calcium imaging in the mice. We recorded a signal and isosbestic channels using excitation wavelengths of 470nm and 415nm, respectively. Before recording, light power at the end of the patch cord was measured to be ~40-60 µw. At the beginning of the behavioral recording, a patch cord was connected to the optical ferrule, and a recording of each channel, in 20Hz, was commenced. In parallel, a video of the behavioral arena using a USB camera (Arducam, B0205 and B0332). To synchronize the video recording with the photometry imaging, a red LED was positioned in the field of view of the video and was turned on upon the start of the photometry imaging. This facilitated the synchronization of the video frames with the calcium imaging data. Photometry data was stored and saved for further analysis.

### Statistical Analysis

All data are presented as mean ± SEM. Data were tested for normal distribution using Shapiro–Wilk's test and Kolmogorov-Smirnov test. Non-parametric test (i.e., Mann–Whitney *U*-test) was chosen when data were found not to be normally distributed.

For IVSA studies, the response rates exhibited during each IVSA phase, schedule (FR1-FR3), and 5F-MDMB-PICA dose tested (1-5 µg/kg/infusion) were analyzed separately by two-way repeated measures (RM) ANOVA with response (i.e., active or inactive lever presses), session and/or schedule as factors, followed by Sidak's multiple comparisons. For RM tests, whenever we could not assume sphericity, a Geisser-Greenhouse correction was carried out by GraphPad Prism 8 software (GraphPad Prism). The session performed under PR schedule was analyzed using unpaired t-test. Considering 5F-MDMB-PICA intake, the data (mean ± SEM of 5F-MDMB-PICA consumption at each dose) were analyzed by ordinary one-way ANOVA followed by Tukey's multiple comparisons post hoc test.

For behavioral experiments, the data were analyzed by using Student's *t*-test when normally distributed and Mann-Whitney test when non normally distributed. Differences were considered significant at $p < 0.05$. Effect sizes were calculated by using Cohen'*d* or Hedges'*g* when sample sizes were equal or not equal, respectively. Post hoc *tests were conducted* only when a significant main effect and/or interaction were detected. All analyses were performed using the GraphPad software package (Prism, version 8; GraphPad, San Diego, California, USA).

For neurochemical studies during odor reactivity test, the effect of odor exposure on DA responses in the mPFC of 5F-MDMB-PICA or Vehicle exposed mice was analyzed by RM two-way ANOVA (treatment x time) followed by Tukey's multiple comparisons post hoc test.

For fiber photometry, analysis was done using scripts written in MATLAB and Python and utilizing known code packages. For video analysis, the mouse position in each video frame was determined using DeepLabCut (12) and Simba (13) to further analyze the mouse behavior in the videos.

In this study, P values $< 0.05$ were considered as statistically significant. Post hoc tests were conducted only when a significant main effect and/or interaction were detected. Statistical analysis was performed with GraphPad Prism 8 (Prism, version 8; GraphPad, San Diego, California, USA) software. Moreover, an effect size study was conducted to provide a clearer understanding of the practical significance of the results obtained from various statistical tests. Different indicators of effect size were used depending on the type of test performed. For Two-way Repeated Measures ANOVA, One-way ANOVA, and F test to compare variances, Eta squared ($\eta^2$) was used to measure the magnitude of the main effects. For the Mixed-effects model (REML), Partial Eta squared ($\eta^2_p$) was employed. For post-hoc comparisons, including Sidak's, Tukey's, and Dunnett's multiple comparisons tests, Cohen's d (d) was calculated to quantify the effect size. In the case of the Mann-Whitney test, the effect size was represented by the correlation coefficient (r). For specific t-tests, R squared ($R^2$) was used to assess the proportion of variance explained by the model. The interpretation of the effect sizes followed standard guidelines. Cohen's d (d): Small: 0.2, Medium: 0.5, Large: 0.8; Eta squared ($\eta^2$) and Partial Eta squared ($\eta p^2$): Small: 0.01, Medium: 0.06, Large: 0.14; Correlation coefficient (r): Small: 0.1, Medium: 0.3, Large: 0.5; R squared ($R^2$): Small: 0.02, Medium: 0.13, Large: 0.26. These measures were used to provide a comprehensive interpretation of the data, highlighting the practical significance of the observed differences beyond the p-values alone.

*REFERENCES*

7. Koolhaas JM, Coppens CM, De Boer SF, Buwalda B, Meerlo P, Timmermans PJA. The Resident-intruder Paradigm: A Standardized Test for Aggression, Violence and Social Stress. JoVE. 2013 Jul 4;(77):4367.
